# Isolation, cloning and expression analysis of two pyruvate orthophosphate dikinase genes of *Chlamydomonas reinhardtii*

**DOI:** 10.1101/2024.05.03.592337

**Authors:** Fadime Demirel, Nicat Cebrailoglu, Faxriyya Mammadova, Taha Tangut, Munevver Aksoy, Gulshan Mammadova, Gulnara Hasanova, Tarlan Mamedov

**Affiliations:** Akdeniz University, Department of Agricultural Biotechnology, Dumlupınar Boulevard 07058 Campus, Antalya, Turkey; Institute of Molecular Biology and Biotechnology, Institute of Molecular Biology and Biotechnologies, The Ministry of Science and Education, Baku, Azerbaijan

**Keywords:** *Chlamydomonas reinhardtii*, pyruvate orthophosphate dikinase (PPDK), gene expression analysis, PPDK activity

## Abstract

The green algae *C. reinhardtii* serves as a useful model for studying photosynthetic cells and has been extensively utilized for investigating various physiological processes. Currently, limited information is available regarding the molecular mechanisms controlling oil accumulation in microalgae. C4 photosynthesis metabolic pathways are essential for high rates of CO_2_ fixation in plants. High rates of photosynthesis are crucial for the biomass accumulation of algae. Surprisingly, C4 pathway enzymes and their regulatory factors have not been studied at the molecular level in any green algae, except for our efforts, which focused on the molecular characterization of phosphoenolpyruvate carboxylase (PEPC) genes (Ppc) in *C. reinhardtii* (Mamedov et al., 2005; Moellering et al., 2007) and phosphoenolpyruvate carboxykinase (Cebarailoglu, 2017). In this study, we isolated and cloned two pyruvate orthophosphate dikinase (PPDK) genes from the green microalga *C. reinhardtii* for the first time and performed expression analysis under different conditions. We demonstrate that both *ppdk* genes encode functional PPDK enzymes in *C. reinhardtii* and that both genes are responsive to changes in carbon dioxide or ammonium concentration during growth. Phylogenetic analysis suggests that *C. reinhardtii* PPDK2 is evolutionarily closer to PPDKs from plants rather than to protozoan and bacterial enzymes. Furthermore, alignment data indicate that the global structure and key amino acid residues involved in catalysis and substrate binding are well conserved in both PPDK enzymes in plants, *C. reinhardtii*, bacteria, and protozoa.

## INTRODUCTION

Recently, microalgae have attracted significant interest worldwide due to their wide range of applications in renewable energy, pharmaceuticals, food, and cosmetic industries. Molecular technologies, including nuclear and organellar (chloroplast, mitochondria) transformation systems, RNA interference, and CRISPR, have been applied to *C. reinhardtii*. These attributes make *C. reinhardtii* an ideal organism for genetic and metabolic engineering to increase oil accumulation in microalgae. *Chlamydomonas reinhardtii* is a unicellular green alga with great potential for biofuel production and has been the subject of extensive research in the field of bioenergy due to its ability to produce lipids (fats) that can be converted into biofuels, primarily biodiesel (Scranton et al., 2015)*. C. reinhardtii* can accumulate a significant amount of lipids, particularly under stress conditions or when subjected to nutrient deprivation (Yang et al., 2015; Choi et al., 2022). These lipids can be used to produce biodiesel, which is a renewable and sustainable alternative to fossil fuels. While *C. reinhardtii* holds promise for biofuel production, there are still challenges to overcome, such as optimizing growth conditions, improving lipid productivity, and developing cost-effective cultivation and harvesting methods. At this point, research strategies need to be developed to make algal biofuels more economically viable and competitive with traditional fossil fuels. Biomass enhancement and metabolic engineering strategies would be essential for optimizing the yield of lipids (oils) that can be converted into biofuels.

Pyruvate orthophosphate dikinase (PPDK; EC 2.7.9.1) is a critical enzyme in the CO_2_ concentrating mechanism (CCM) known as the C4 pathway (Ashton et al., 1990). PPDK catalyzes the conversion of pyruvate, adenosine triphosphate (ATP) and inorganic phosphate (Pi) to phosphoenolpyruvate (PEP), adenosine monophosphate (AMP), and pyrophosphate (PPi) in a wide variety of organisms, ranging from bacteria to eukaryotes (Hatch & Slack, 1968; Taylor et al., 2010).

In C4 plants, PPDK plays a crucial role in the C4 photosynthesis pathway, which is an adaptation to overcome photorespiration and increase carbon fixation efficiency, especially in hot and arid environments (Roeske et al., 1988; Roeske et al., 1989; Smith et al., 1994a; Smith et al., 1994b). In C4 plants, PPDK is localized in mesophyll cells, where RuBisCO operates without exposure to atmospheric oxygen, reducing photorespiration.

Several studies have shown that PPDK is critically controlled by light in C4 plants (Chastain 2003; Edwars et al., 1985; Furbank 1995; Roeske et al., 1988; Roeske et al., 1989; Smith et al., 1994a; Smith et al., 1994b). PPDK is also found in C3 (wheat, rice, tobacco, etc.) plants, but its role and regulation differ between these two types of plants due to the differences in their photosynthetic pathways. In the C3 pathway, carbon dioxide is initially fixed into a three-carbon compound, 3-phosphoglycerate (3-PGA), via ribulose-1,5-bisphosphate carboxylase/oxygenase (RuBisCO) (Biswal et al., 2018). In C3 plants its main role is in other metabolic processes, such as the conversion of pyruvate to phosphoenolpyruvate (PEP) and vice versa in glycolysis and gluconeogenesis (Daloso et al., 2017).

In summary, while PPDK is present in both C3 and C4 plants, their primary roles and locations are different. In C3 plants, it has a role in general metabolism, whereas, in C4 plants, it is a key enzyme in the C4 carbon fixation pathway, helping to increase photosynthetic efficiency by reducing photorespiration and enhancing carbon dioxide concentration in bundle sheath cells. This difference in PPDK function is one of the key distinctions between C3 and C4 photosynthesis. Generally, green algae have been considered C3 plants, similar to most terrestrial plants. However, evidence suggests that certain green algae species have evolved mechanisms that resemble C4 photosynthesis in certain aspects. *Ulva prolifera*, a typical green tide-forming alga, has been shown to have both the C3 and C4 photosynthetic pathways (Xu et al., 2012).

*C. reinhartii* are known to generally perform C3 photosynthesis, still possible that *C. reinhardtii* might exhibit some characteristics of C4-like photosynthesis under certain conditions. Notably, C4 pathway enzymes and their regulatory factors have not been studied at the molecular level in any green algae, except our previous efforts, which were focused on the molecular characterization of PEPC genes (*Ppc)* in *C. reinhardii* (Mamedov et al., 2005; Moellering et al., 2007) and carboxykinase genes in *C. reinhardtii* (Cebrailoğlu, 2017). Thus, given the complete lack of molecular and biochemical insight into the enzymes of C4-metabolism in any green algae, our research has been focused on investigating one of the key enzymes of C4-metabolism enzymes, PPDK in the photosynthetic model organism *C. reinhardtii*. Although PPDK has not been studied in *C. reinhardtii*, however, there are few studies on this enzyme in some algae (Huang et al., 2023). In plants and some algae isoforms of PPDK proteins are found in both the chloroplast and cytosol (Sheen, 1991; Parsley & Hibberd, 2006). PPDK is encoded by nuclear DNA and the gene is transcribed from two different initiation sites under the control of two promoters: the larger transcript produces the chloroplastic PPDK, which includes a transit peptide, and the smaller one produces the cytosolic PPDK (Parsley & Hibberd, 2006). Notably, earlier studies demonstrated that maize has been found to contain two PPDK genes: one that encodes a cytosolic isoform, and a second PPDK gene that either generates long transcripts encoding chloroplastic PPDK or shorter transcripts encoding a cytosolic isoform (Sheen, 1991). PPDK genes have been isolated from rice and the genome was found to contain two *ppdk*, namely *osppdka* and *osppdkb* (Moons et al., 1998). Parsley and Hibberd (2006) reported that the single gene encoding PPDK in *Arabidopsis thaliana* possesses two promoters. In this work, full-length cDNA–gfp fusions were produced to investigate the subcellular localization of the two PPDK enzymes. While the cDNA derived from the longer transcript and GFP led to protein accumulating in chloroplasts, the PPDK-GFP fusion derived from the shorter transcript appeared to localize to the cytosol of the cell (Parsley & Hibberd, 2006).

The unicellular algae *C. reinhardtii* is an important model organism for studying essential biological processes and also biotechnological applications (Harris, 2001; Merchant et al., 2007; Salomé & Merchant, 2019; Sasso et al., 2018). Two isoforms of pyruvate phosphate dikinase, PPDK1 and PPDK2 have been reported in *C. reinhardtii* (Boyle et al., 2017; Huang et al., 2008). Wakao et al. (2021) sequenced the genomes of 660 *C. reinhardtii* acetate-requiring mutants, part of a larger photosynthesis mutant collection. PPDK2 was predicted to be chloroplastic (Terashima & Hippler, 2011). According to Phytozome version 13, while PPDK1 (Cre10.g424750) is probably cytosolic, PPDK2 (Cre17.g734548_4532) is predicted to be localized in the chloroplast. Similarly, using the CELLO program, it was predicted that PPDK1 is in the cytosol and PPDK2 is localized in the chloroplast. However, no direct studies are confirming that these proteins are present in the cytosol and/or chloroplast in *C. reinhardtii*, so the functions of these proteins are not fully understood. It should be noted that chloroplast PPDK has been characterized to play a key role in the regeneration of phosphoenolpyruvate in the chloroplasts of mesophyll cells to maintain the C4 cycle (Astley et al., 2011; Hatch, 1987).

In a previous study with rice seeds, it has been claimed that the cytosolic PPDK protein plays a role in regulating the flux of carbon into starch and fatty acids (Kang et al., 2005). In other studies, cytosolic PPDK is also proposed to be involved in amino acid interconversions and regulation of the equilibrium between starch and storage protein accumulation in seeds (Astley et al., 2011; Chastain, 2006; Méchin et al., 2007) .

In this study, for the first time, we reported the isolation and cloning, overexpression in *E. coli* and gene expression analysis of two *ppdk* genes from green algae *C. reinhardtii.* We demonstrated that both *ppdk1* and *ppdk2* genes encode functionally active PPDK enzymes in *C. reinhardtii.* We also demonstrate that two *PPDK* genes are C- and N-responsive genes and up-/down-regulated by changes in [CO_2_] or [NH_4_^+^] during growth.

## Results and Discussion

### Isolation and cloning of *C. reinhardtii PPDK* genes and construction of expression vector

Based on similarity and previous studies, on homology to *Flaveria pringlei* and TargetP predictions (May et al., 2009; Emanuelsson et al., 2000), *ppdk* genes (*ppdk1*, and *ppdk2*) have been predicted and reported in *C. reinhardtii,* of which PPDK2 is most likely chloroplastic (May et al., 2009; Emanuelsson et al., 2000). The full length of the *ppdk1* and *ppdk2* genes from *C. reinhardtii* cDNA was isolated and cloned using reverse transcription-polymerase chain reaction (RT-PCR) with specific primers listed in Table 1. Agarose gel electrophoresis of the PCR products showed that 2682-bp of *ppdk1* and 2904-bp of *ppdk2* genes at the expected sizes were successfully isolated and cloned (Figure 1).

**Figure 1.**
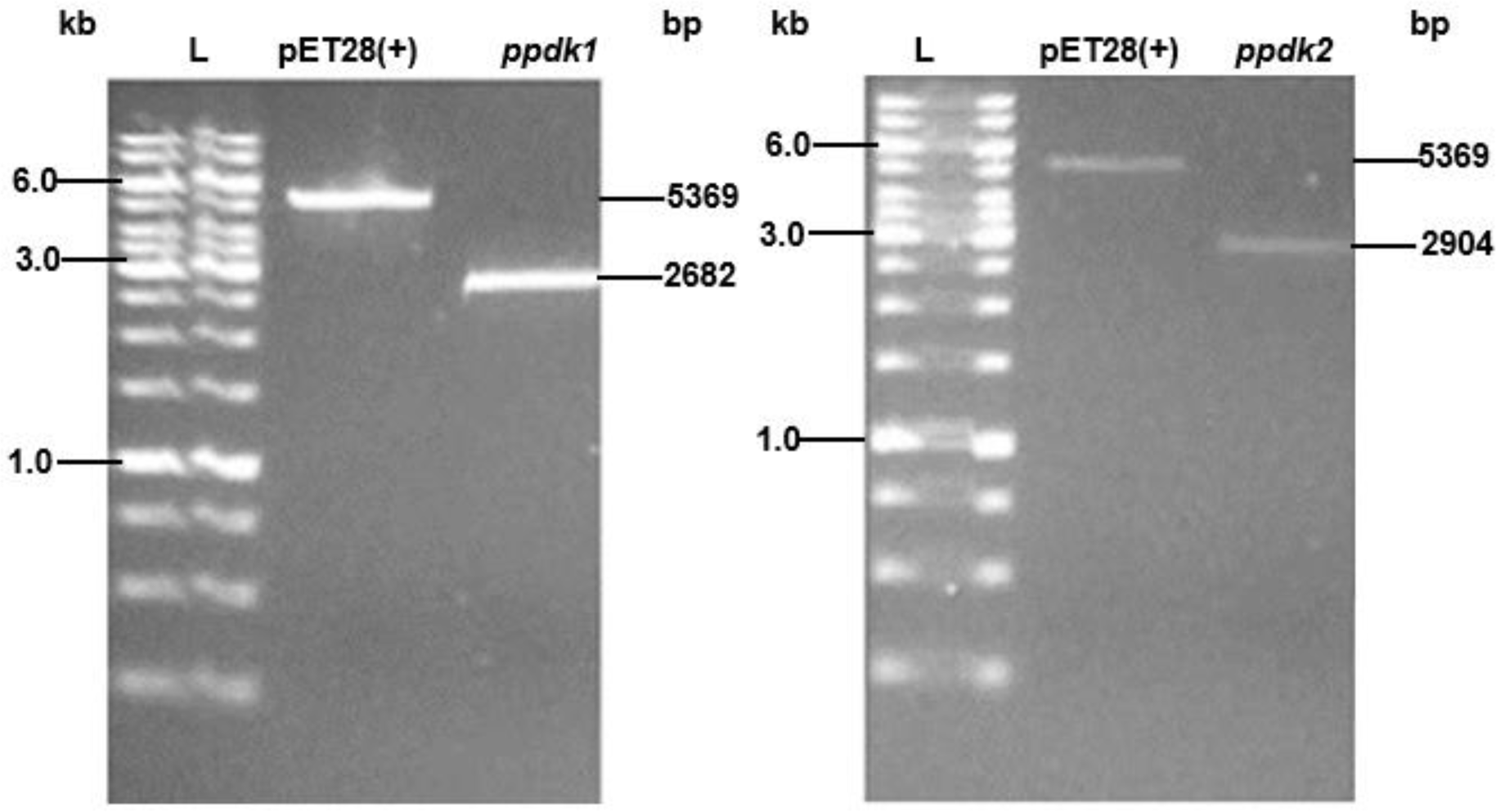
Agarose gel electrophoresis of *Nde I* and *Xho I* digested pET-28a (+)-ppdk1 or pET-28a(+)-ppdk2 plasmids. Digestion releases pET-28a (+) vector (5369 bp), *ppdk1* (2682 bp) and *ppdk2* (2904 bp ) genes of *C. reinhardtii*. L: GeneRuler 1kb DNA Ladder (Thermo Scientific).

**Table 1:**
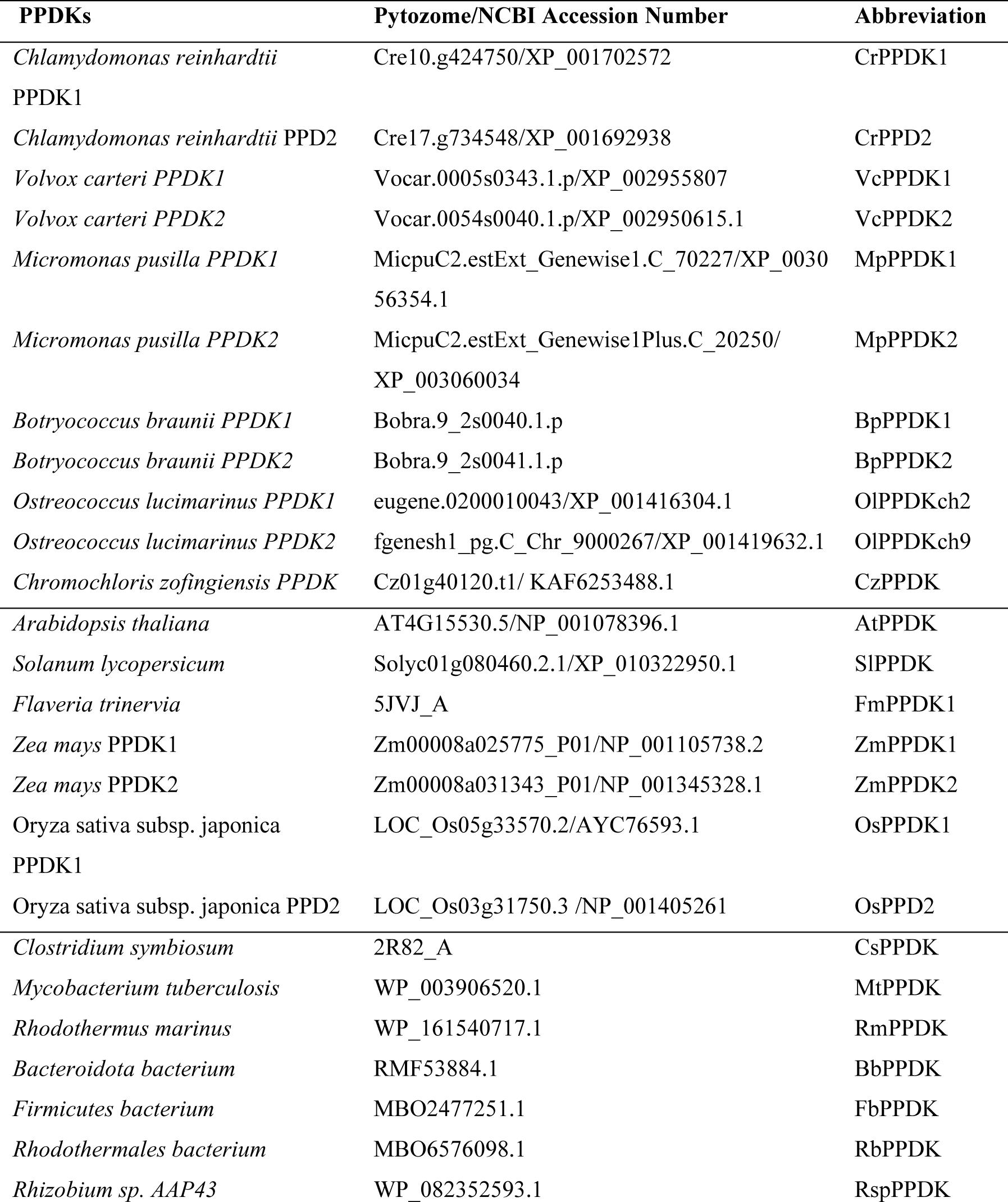
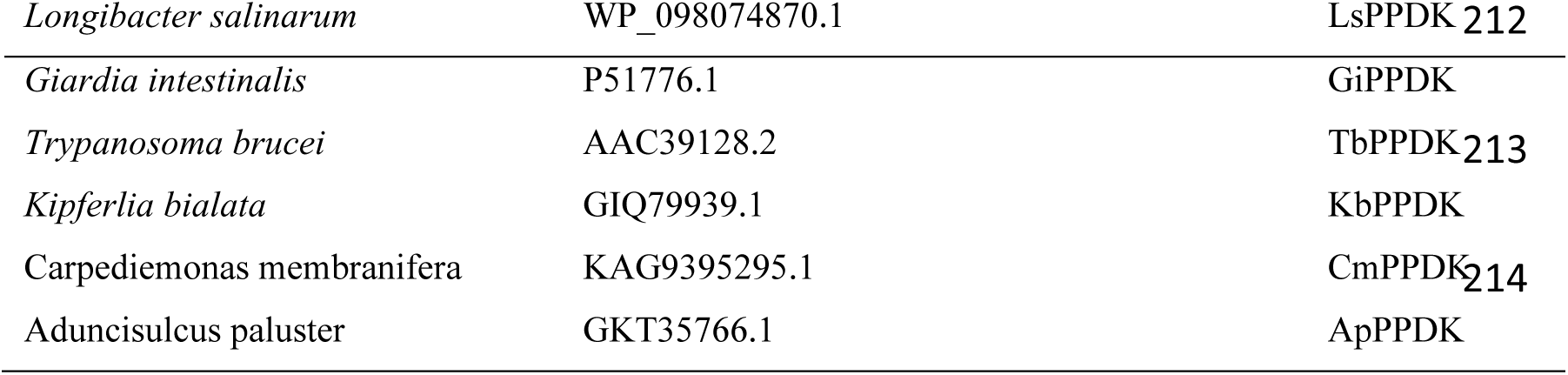
Pytozome/NCBI accession numbers of various PPDKs.

### Overexpression of PPDK1 and PPDK2 in *E. coli*

We overexpressed *C. reinhardtii* ppdk1 and ppdk2 genes in *E. coli* to determine if they encode functional PPDK enzymes in *C. reinhardtii*. The full-length *ppdk1* and *ppdk2* genes were cloned into *E. coli* expression vector pET28a(+) downstream of the T7 promoter for IPTG-inducible transcription by the T7 RNA polymerase and to express the PPDK1 and PPDK2 proteins with 6x histidine tag (Figure 2a, b). *E. coli* cells transformed with pET28a(+)-*ppdk* genes were induced for 3h with 0.5 mM isopropylβ-D-thiogalactoside (IPTG) at 25 °C. *E. coli* cells, which were transformed with an empty pET28a (+) vector and induced with 0.5 mM IPTG were used as a negative control. As can be seen from Figure 2, the induction of approximately 100 kDa proteins was observed. The PPDK protein was not observed in the negative control, i.e. the component *E. coli* cells transformed with pET28a (+) vector The predicted molecular mass (MM) of the PPDK1 and PPDK2 proteins are approximately 96.689 kDa and 104.451 kDa, and the predicted isoelectric points (pI) are 5.90 and 5.48, respectively, based on calculations using Prot-Param tool (https://web.expasy.org/protparam/). MM calculated using the Prot-Param tool are consistent with MM of recombinant PPDK1 or PPDK2 proteins (Figure 2c). The expressed recombinant PPDK proteins were also found to be highly soluble, evidenced based on the results of western blot analysis. Enzymatic activities of PPDK enzymes were determined in total cell extracts. Both PPDK1 and PPDK2 recombinant enzymes were found to be enzymatically active; PPDK2 had more activity compared to PPDK1 (Figure 2d). Thus, based on these analyses, we conclude that ppdk genes encode functionally active enzymes in *C. reinhardtii*.

**Figure 2.**
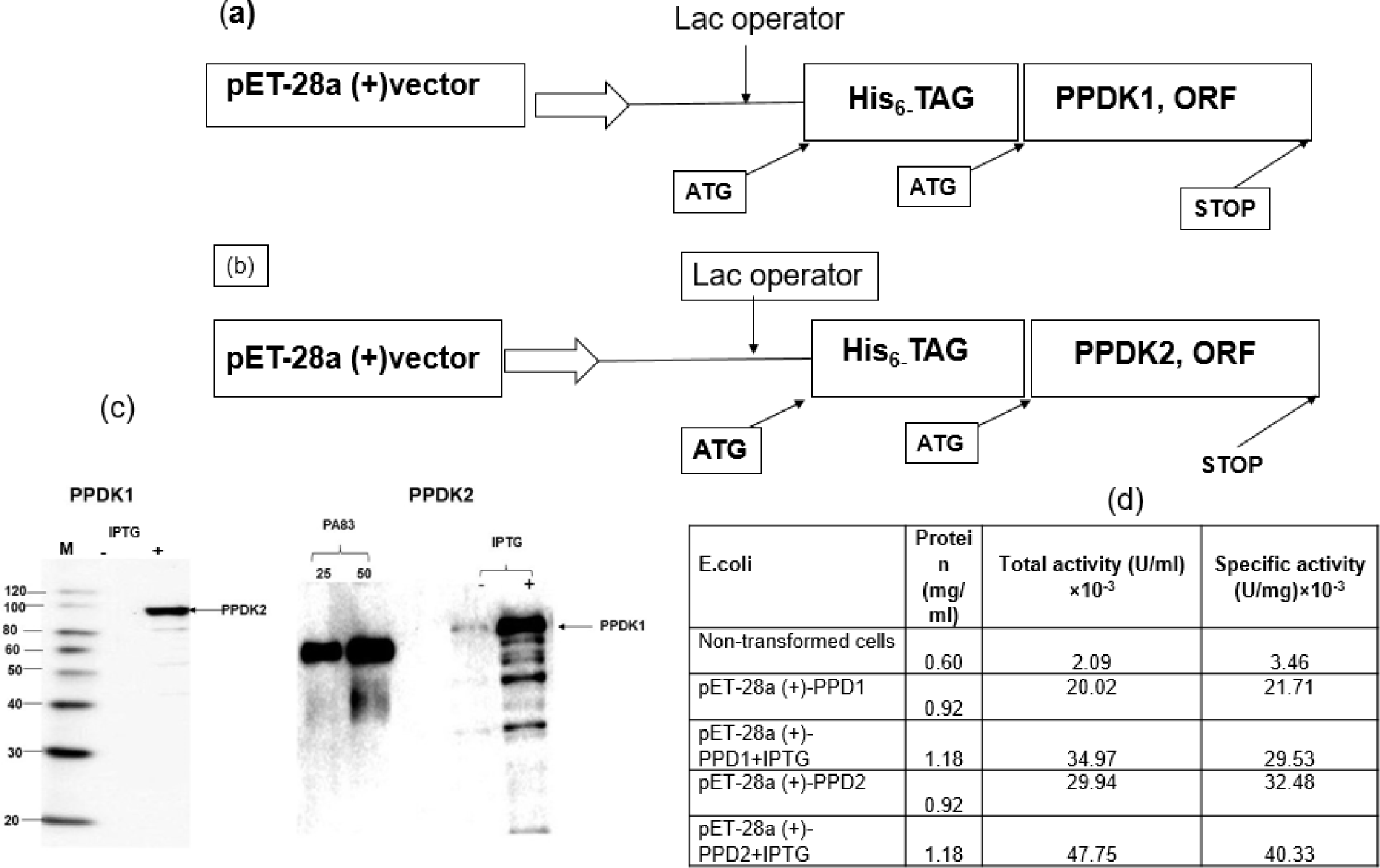
Recombinant expression of PPDK1 and PPD2 in *E. coli* and subsequent activity assessment. (a, b) schematic representation of pET-28a(+) constructs expressing*ppdk1* and *ppdk1* genes from *C. reinhardtii*. (c) Immunoblot analysis of PPDK1 and PPD2 recombinant proteins expressed in *E. coli* strain BL21(DE3). M: MagicMark XP Western Protein Standard (ThermoFisher Scientific). (d) activity assessment of *E. coli* extract expressing *ppdk1* and *ppdk1* genes. Protein extract of non-transformed *E. coli* (control), - IPTG: Protein extract of transformed *E. coli* with no IPTG added; + IPTG: Protein extract of transformed *E. coli* with IPTG added. pp-PA83: plant produced PA83 of *Bacillus anthracis,* developed and purified in our laboratory (Mamedov et al., 2017) and used as a protein marker with a molecular mass of about ∼80 kDa.

### Amino acid sequence alignments

Amino acid sequence alignment of *C. reinhardtii* PPDK1 and PPDK2 are shown in Figure 3. It was observed that the GGMTSHAAVVA motif, which is conserved in the PPDK enzymes, is also conserved in *C. reinhardtii* PPDK1 and PPDK2. The protein sequences of *C. reinhardtii* PPDK1 and PPDK2 share 55.4% identity (Demirel, 2023).

**Figure 3.**
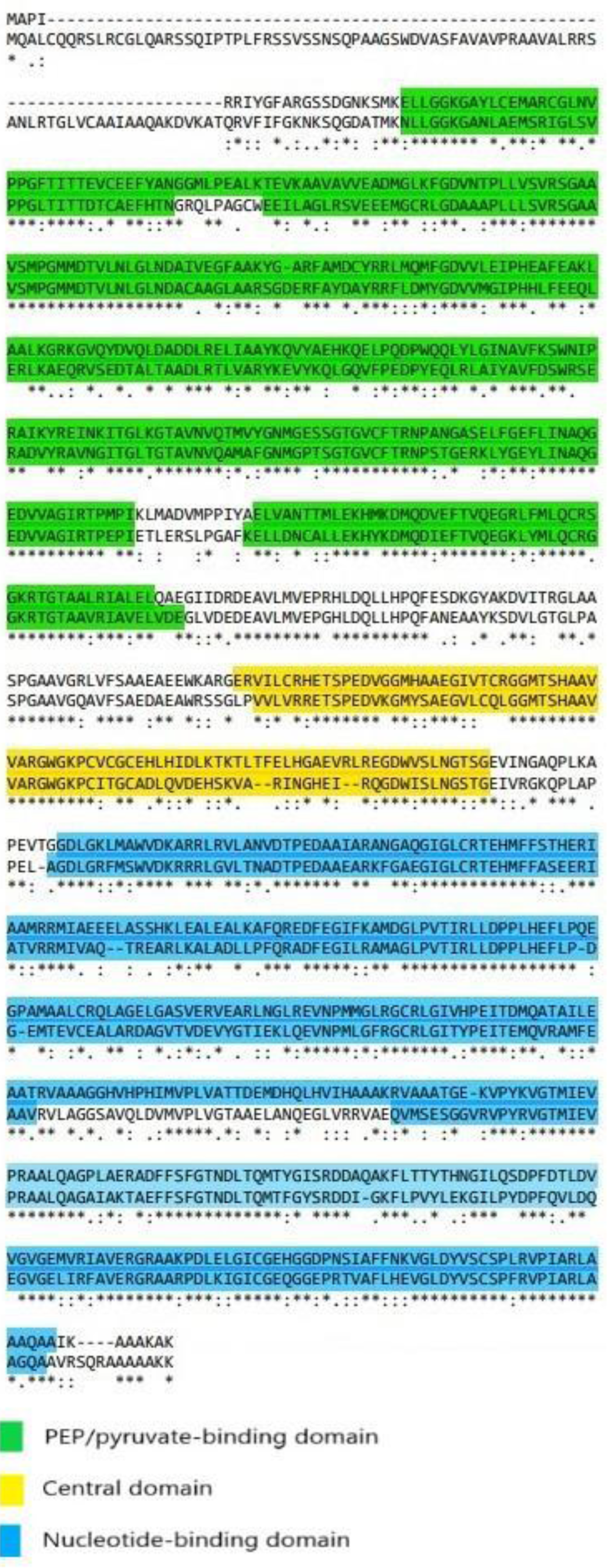
Amino acid sequence alignment of *C. reinhardtii* PPDK1 and PPDK2 proteins.

The results of multiple amino acid sequence alignment showed that PPDK1 and PPDK2 of *C. reinhardtii* share significant homology with the PPDK proteins from algae and plants (Figure 4), such as *V. carteri*, *A. thaliana, Z. mays etc.* In addition, PPDK1 and PPDK2 of *C. reinhardtii* consist of three independently folded biochemically defined functional domains: nucleotide-binding domain (in the N-terminal domain), central domain, and Pep/pyruvate-binding domain (in the C-terminal domain) (Herzberg et al., 1996). The central domain is where the catalytic Histidine is located in the GGMTSHAAVVA motif (Figure 3). This region includes histidine and threonine residues, which are presumed to serve as the catalytic site and the site of phosphorylation and dephosphorylation that regulate the reversible activation of this enzyme (Agarie et al., 1997; Edwards et al., 1982; Imaizumi et al., 1997).

**Figure 4.**
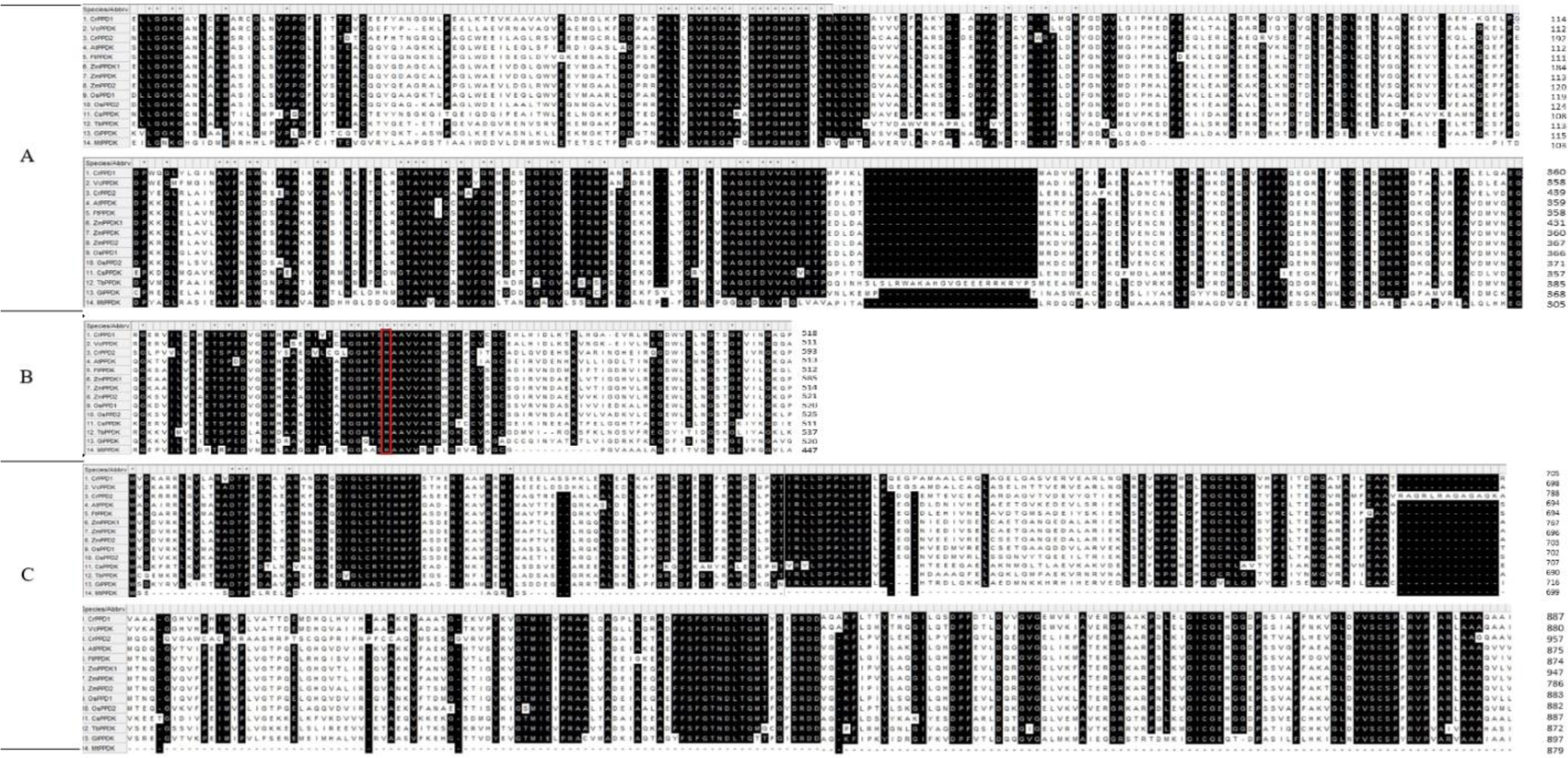
Selected amino acid sequence alignments of two PPDK1 and PPDK2 amino acid sequences from *C. reinhardtii* and representative vascular plants, humans, algae, *Cyanobacteria,* and prokaryotic species. Accession numbers are given in Table 1. Conserved amino acid residues are shown in black. The catalytic histidine residue is marked with a red color.

To investigate the domains of the PPDK, PPDK proteins were selected from various organisms such as algae, plants, and bacteria (Figure 5). Our results show that, with the exception of the PPDK from *Mycobacterium tuberculosis*, the PPDKs of all other organisms are composed of three domains. Notably, *C. reinhardtii*, *Volvox carteri*, *Micromonas pusilla, Oryza sativa and Zea mays* have two PPDK isoforms.

**Figure 5.**
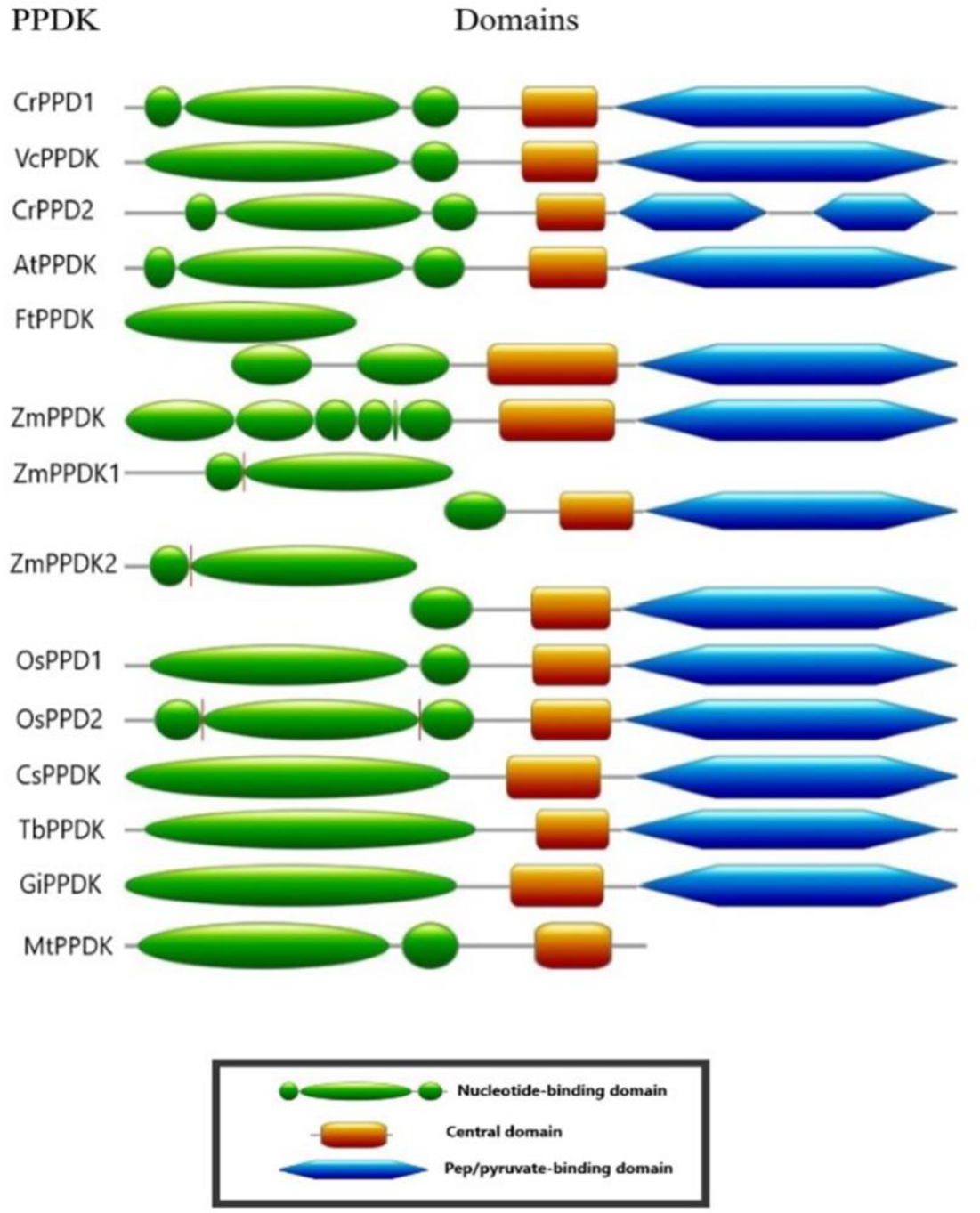
Domains of PPDK proteins.

Several studies have shown that PPDK is critically controlled by light in plants with the C4 pathway. It should be noted that there is a completely conserved amino acid sequence (GGMTSHAAVVA) in the PPDK protein in C4 and C3 plants. The conserved GGMTSHAAVVA sequence was also found in PPDK1 and PPDK2 sequences in *C. reinhardtii*. The conservation of this sequence around the regulatory site of rice, maize and *C. reinhardtii* PPDKs suggests that the regulatory mechanisms of PPDK enzymes could be similar in C3 and C4 plants and also in *C. reinhardtii*.

### Phylogenetic analysis

PPDK1 and PPDK2 sequences of *C. reinhardtii* were compared to PPDK sequences of *V. carterii*, *A. thaliana*, *O. sativa, Z. mays*, *C. symbiosum*, *F. trinervia*, *T. brucei*, *G. intestinalis* and *M. tuberculosis* (Figure 6). According to our analysis, *C. reinhardtii* PPDK2 sequence was more closely related to plant PPDKs while *C. reinhardtii* PPDK1 was closer to its closest relative *V. carterii.* As can be seen from Figure 6, *C. reinhardtii* PPDK1 and PPDK2 sequences were grouped under different sub-clusters.

**Figure 6.**
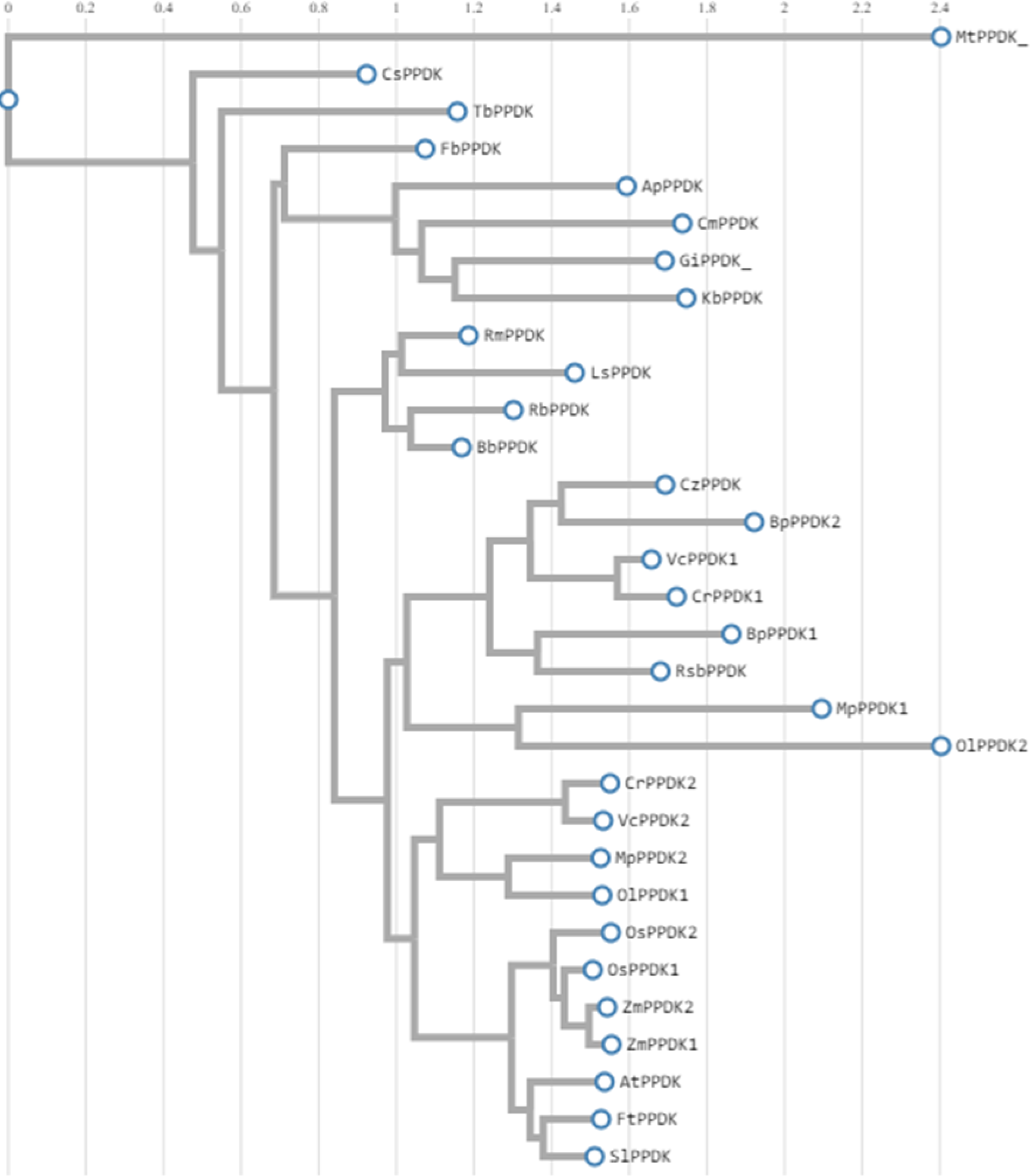
Phylogenetic relationship of *C. reinharditti* PPDK1 and PPDK2 proteins.

### Enzyme Activity of *C. reinhardtii* Cells grown **under** high and low CO_2_ conditions

When high CO2-grown *C. reihardtii* cells switched to air for 0.5 h, 1 h and 3 h, the specific activity of the PPDK enzyme decreased in 7.27, 4.44 and 3.74 -fold, respectively. When air-grown cells switched to high CO_2_ (5 %) for 0.5 h, 1 h and 3 h, the specific activity increased at 1.07, 4.11 and 3.9, respectively (Table 2). These results support a photosynthetic role of both PPDK1 and PPDK2 in *C. reinhardtii*.

**Table 2:** PPDK activities of *C. reinhardtii* cells grown in high CO_2_ and switched to air (0.5, 1 and 3 hours) or grown in ambient air (low CO_2_) and then switched to air (0.5, 1 and 3 hours). The data presented is the meaning of at least 3 biological replicates. One unit of activity (U) is equal to the conversion of one μmol of substrate per minute. Specific activity is the activity per mg cell extract protein.

|  | Protein (mg/ml) | Total activity (U) | Specific activity (U/mg) | Specific activity (%) | Fold for specific activity |
| --- | --- | --- | --- | --- | --- |
| High CO<sub>2</sub> grown cells | 1.40 | 27.28 | 19.48 | 100 | 1 |
| High CO<sub>2</sub> switched to air 0.5 h | 1.302 | 3.486 | 2.68 | 13.49 | 7.27 Fold decrease |
| High CO<sub>2</sub> switched to air 1 h | 0.779 | 3.422 | 4.39 | 22.14 | 4.44 Fold decrease |
| High CO<sub>2</sub> switched to air 3 h | 0.885 | 4.610 | 5.21 | 26.73 | 3.74 Fold decrease |
| Air grown cells | 1.325 | 4.243 | 3.20 | 100 | 1 |
| Air grown cells switched to high CO<sub>2</sub>, 0.5 h | 1.180 | 40.69 | 3.44 | 107.43 | 1.07 Fold increase |
| Air grown cells<br>switched to high<br>CO<sub>2</sub>,<br>1 h | 1.063 | 13.20 | 13.16 | 410.99 | 4.11<br>Fold increase |
| Air grown cells<br>switched to high<br>CO<sub>2</sub>,<br>3 h | 1.123 | 14.01 | 12.47 | 389.57 | 3.90<br>Fold increase |

### Enzyme Activity of *C. reinhardtii* cells grown under high and low NH^+^ concentrations

When *C. reihardtii* cells were grown in a TAP medium (2.1 mM NH^+^_4_) and switched to a TAP medium containing 3.97 mM or 9.58 mM NH^+^ , the specific activity of the PPDK enzyme increased by 1.22 or 1.5-fold, respectively (Table 3). The increase in specific activity of PPDK was also accompanied by an increase in soluble total protein in the culture.

**Table 3.** PPDK activities of *C. reinhardtii* cells grown under different NH^+^_4_ concentrations in TAP-medium

| NH<sub>4</sub><sup>+</sup><br>concentration<br>(mM) | Protein<br>concentration<br>(mg/ml) | Total enzyme<br>activity<br>(U/ml) | Specific activity<br>(U/mg) | Fold increase<br>for a specific<br>activity |
| --- | --- | --- | --- | --- |
| 9.58 | 5.46 | 127.8 | 23.41 | 1.5 |
| 3.97 | 3.84 | 72.9 | 18.98 | 1.22 |
| 2.1 | 3.10 | 48.1 | 15.52 | 1.0 |

### Expression analysis of two *PPDK* genes in *C. reinhartii g*rown under different CO_2_ concentrations

We performed expression analysis of *ppdk1* and *ppdk2* genes of *C. reinhardtii* cells grown in high CO_2_ (∼5%) and low CO_2_ (ambient air, ∼0.04% CO_2_) in HS medium Acetate is a known organic carbon source for *C. reinhardtii*, its uptake requires ATP and its catabolism produces NADPH (Johnson & Alric, 2012; Plancke et al., 2014). The expression of both *ppdk1* and *ppdk2* genes of *C. reinhardtii* increased in cultures transferred from air to high CO_2_ concentrations. Expression levels of *ppdk* genes increased after 5 h of high CO_2_ supply. After 24 h, the expression of these genes increased approximately 60-fold compared to the initial levels of the *ppdk1* gene, while ppdk2 increased 10-fold (Figure 7). These gene expression analyses well correlate with enzyme activity and support the photosynthetic role of both PPDK1 and PPDK2 in *C. reinhardtii.* Notably, the reference gene, carbonic anhydrase gene (*CAH1*) was down-regulated when air-grown cells transferred to high CO_2_ as expected, which is consistent with the previous studies (Mamedov et al., 2005).

**Figure 7.**
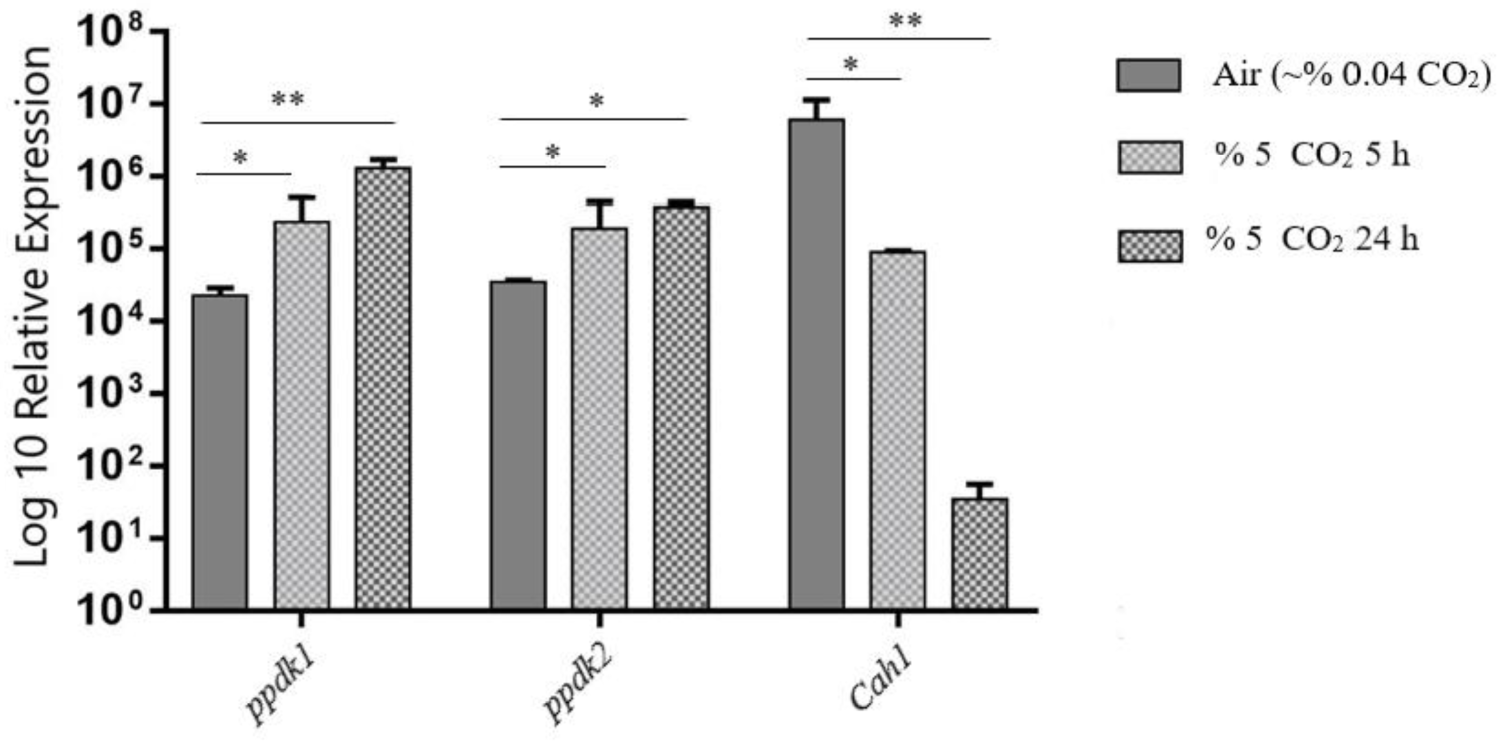
Reverse transcription quantitative real-time PCR analysis of *PPDK1*and *PPDK2* genes and CO_2_-responsive control transcript (*Cah1*) (*n* = 3, t = 2) in *C. reinhardtii* cells grown in different levels of CO_2_.

### Expression analysis of two *PPDK* genes under different NH^+^_4_ concentrations

Expression analysis of two *ppdk* genes and the ammonium transporter (*Amt4*), as inorganic-N responsive reference gene, were performed by qRT-PCR at low (0.5 mM) and high (10 mM) NH^+^_4_ conditions. The N-responsive *Amt4* gene was shown to be upregulated at N-deficient or low (0.5 mM) N concentrations (Mamedov et al. 2005; Mamedov et al., 2024). The *Amt4* gene (AY542491) used in this study, is one of four *Amt* genes in *C. reinhardtii*, which was shown to encode a putative gas channel for the uncharged NH_3_ species (Soupene et al., 2004). As can be seen from Figure 8, NH^+4^ concentrations of the *Amt4* gene, is upregulated at low ammonium levels (0.5 mM), which is consistent with our previous studies (Mamedov et al., 2005; Mamedov et al., 2024). When cells grown at 10 mM ammonium were switched to 0.5 or 1.0 mM, no significant change was observed in *PPDK1* gene expression. However, at the same condition, when cells grown in high ammonium (10 mM) switched to low ammonium (0.5 or 1.0 mM), the *ppdk2* gene was down-regulated and then when cells switched back to high ammonium (10 mM) for 5 and 24 h, the *ppdk2* gene was up-regulated. As shown above, when *C. reihardtii* cells grown in TAP medium (2.1 mM ammonium) were switched to TAP medium containing 3.97 mM or 9.58 mM ammonium, the specific activity of the PPDK enzyme increased by 1.22 or 1.5 times, respectively, despite an increase in total soluble protein (Table 3). Since the *ppdk2* gene is upregulated at high ammonium these data suggest that the increase in specific activity of the PPDK enzyme in a high ammonium TAP medium is probably due to the increase of the PPDK2 enzyme. Collectively all these results support that the *ppdk2* gene is an N-responsive gene and is up- and down-regulated by changing the ammonium concentration in the growth medium and support a non-photosynthetic role of PPDK2 in *C. reinhardtii* (Edwards et al., 2001; Mamedov et al., 2005).

**Figure 8.**
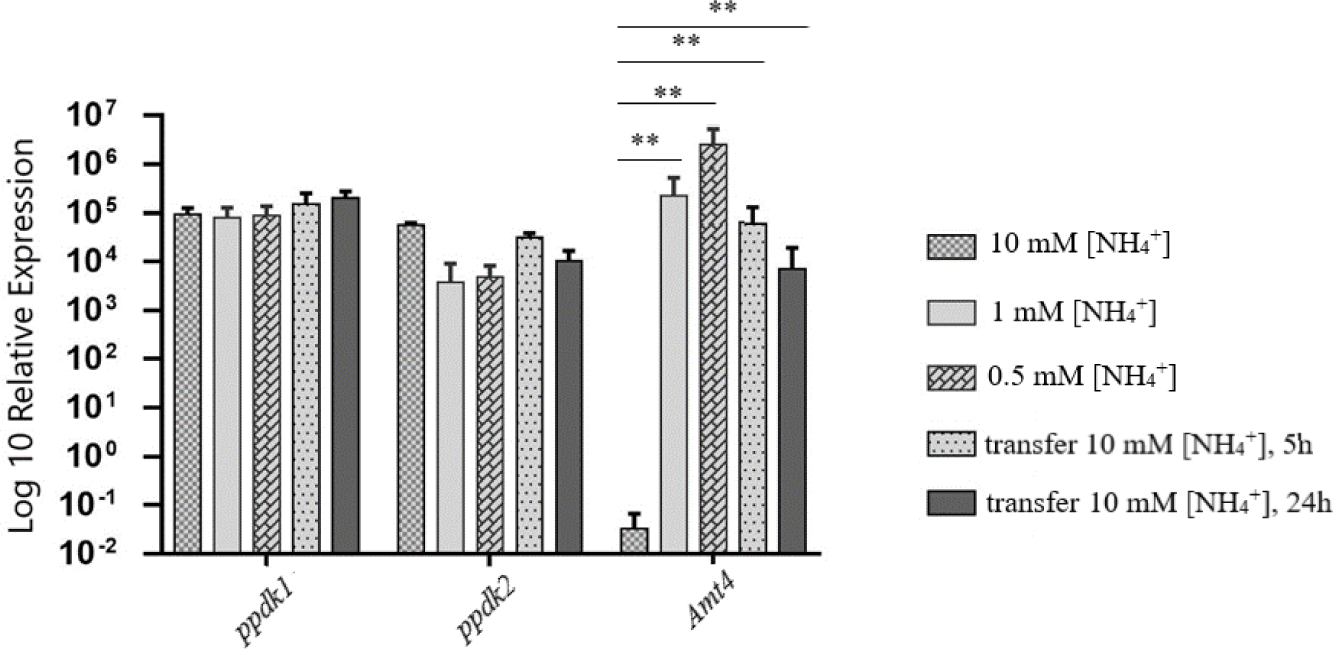
Expression levels of *PPDK1*, *PPDK2* and *AMT4* of *C. reinhardtii* (*n* = 3, t = 2), grown in high/low levels of ammonium.

## MATERIAL AND METHOD

### Strains and growth conditions

*C. reinhardtii* wild type (wt) strain CC-125 mt^+^ was obtained from Chlamydomonas Genetics Center, Duke University. *C. reinhardtii* strain was cultured at room temperature in a Tris-acetate-phosphate medium (Gorman & Levine, 1965; Siaut et al., 2011) or at high salt (HS) medium as described previously (Harris, 1989; Mamedov et al. 2001, 2005, 2024) by shaking at 120 rpm under continuous light (∼200 photons m^-2^ s ^-1^). Cultures were grown until 2-4 × 10^6^ cells/ml density and cells were separated by centrifugation at 5000 rpm for 15 min. Then the pellets were ground in a mortar containing liquid N_2_ and powders were stored at -80 °C for later analysis.

HS medium was used for RT-qPCR analyses under varying levels of ammonium or CO_2_. Ammonium concentrations used were 0.5, 1.0 or 10.0 mM. CO_2_ conditions were adjusted by bubbling with air enriched with 5% CO_2_ or ordinary air alone. To achieve high levels of CO_2_ concentrations, *C. reinhardii* cultures were bubbled continuously with air enriched with CO_2_ to 5% by volume. To grow *C. reinhardtii* in low CO_2_ conditions, cultures were bubbled with ordinary air (∼0.04% CO_2_) in the same medium.

### RNA isolation and cDNA synthesis

Total RNA was isolated using the TRIzol reagent (Ambion, USA) and treated with DNase I (Thermo Scientific) (Rio et al., 2010; Mamedov et al., 2024). RNA quality and concentration were determined using ethidium bromide (EB) stained agarose gel electrophoresis and BioDrop (Indolab, Utama), respectively. RNA samples were stored at −80 °C until needed for the cDNA synthesis. First-strand cDNA synthesis was performed using One Taq RT-PCR kit (New England BioLabs) and 1μg of total RNA was used according to the manufacturer’s instructions.

### Construction of PPDK expression vector and over-expression in *E. coli*

The plasmid pET-28a (+) (Novagen, Darmstadt, Germany) was employed for the recombinant expression of *C. reinhardtii* PPDK1 and PPDK2 in *E. coli*. Primers used for the cloning of the genes are given in Table 4. cDNAs were used as templates for the amplification of coding sequences of *ppdk2* and *ppdk2* genes. For amplification of *ppdk1*, primers PPDK1 F1 and PPDK1 R1 were used. For amplification of *ppdk2*, primers PPDK2 F1 and PPDK2 R1 were used. PCR was performed using NEB Q5 High Fidelity PCR Kit. Amplifications were performed at 98°C for 30 s, followed by 25–35 cycles of 98°C for 10 s, 50-72°C for 30 s and 72°C for 30 s, and a final extension at 72°C for 2 min. PCR products were separated on a 1% agarose gel in TAE buffer. Ethidium bromide was also added to the agarose gel, and DNA was visualized with the UV imager MiniLumi system (DNR Bio-imaging Systems). After amplification, the PCR product was purified with DNA Clean and Concentrator Kit (Zymo Research, USA). The amplified PCR fragments were then cloned into pGEM-T Easy vector (Promega) and confirmed by sequencing. After confirmation, primers containing *NdeI* and *XhoI* sites (Table 4) were used to amplify the genes to clone into the pET-28a(+) vector.

**Table 4.**
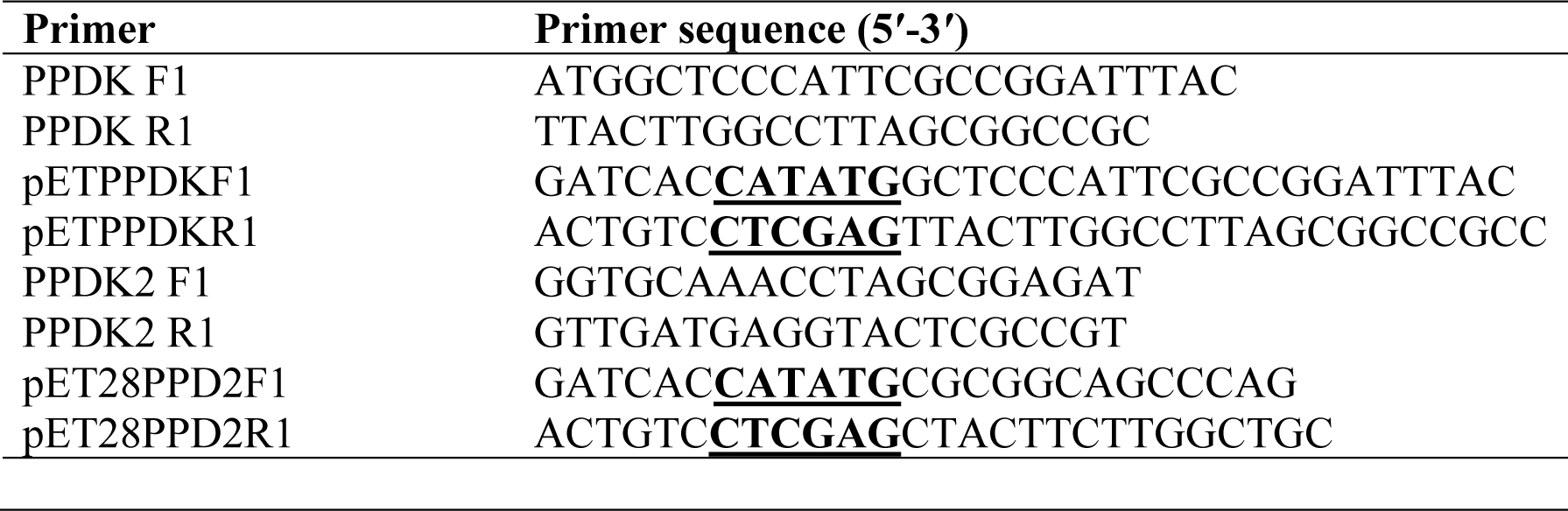
Sequences of primers used for cloning of *ppdk1* and *ppdk2* genes into pET28a(+) bacterial protein expression vector. The bold underlined nucleotide bases indicate *NdeI* and *XhoI* restriction enzyme digestion sites.

For cloning into the pET-28a(+) vector, the vector and the PCR products were digested by *NdeI* and *XhoI* for 10 min at 37°C and then were inactivated at 65°C for 5 min. The digested products were gel purified and ligated for 20 min at room temperature using Quick Ligase enzyme (NEB, DH5a competent cells using the heat shock method.

Plasmids were isolated using Zyppy Plasmid Miniprep Kit (Zymo Research, USA) and confirmed with agarose gel electrophoreses. Confirmed plasmids were transformed into the *E. coli* BL21 (DE3) cells for expression of PPDK1 and PPDK2 proteins. Cultures of *E. coli* strain BL21(DE3) were grown at 37°C in 100 mL of LB medium containing 30 μg/mL kanamycin to an OD_600_ of 358 ∼0.6. Recombinant protein expression was induced using 1 mM isopropyl-b-D-thiogalactoside 359 (IPTG) at 25°C for 3 hours. Expression without IPTG was also performed. The cells were harvested at 4°C by centrifugation at 2000 g for 15 min and re-suspended in 20 mM sodium phosphate extraction and binding buffer containing 10 mM imidazole and 300 mM sodium chloride (pH 7.4). Cells were lysed at 4°C by sonicating (Mamedov et al., 2005) and purified using HisPur*™* Ni-NTA resin ((Thermo Fisher Scientific, Cat. No. 8822), as described recently (Mamedov et al., 2023). The fractions possessing high PPDK activity were combined, desalted and used for PPDK activity assays.

### PPDK activity assay

Samples of 150-200 mg liquid N_2_-frozen *C. reinhardtii* or *E. coli* cells were homogenized in an ice-cold extraction medium. The extraction medium contained 50 mM Tris-HCl (pH 7.4), 10 mM DDT, 10 mM MgSO_4_, 1 mM EDTA, 2 mM KH_2_PO_4_, and 10 mM 2-mercaptoethanol. PPDK activities were determined spectrophotometrically by following NADH oxidation at A_340_ at 25°C. An extinction coefficient of 6.22 AU mmol^−1^ L cm^−1^ was used for NADH. The reaction medium contained 25 mM Hepes-KOH, pH 7.4, 6 mM MgSO_4_, 10 mM DTT, 25 mM NH_4_Cl, 0.25 mM NADH and 2 U/ml of LDH (L-Lactic Dehydrogenase from bovine heart); with saturating concentration of the enzyme’s substrates; 1 mM PEP, 0.5 mM AMP, 1 mM PPi (Bringaud et al., 1998; Fukayama et al., 2001; South & Reeves, 1975). All enzymes were assayed in a final volume of 200 μL using a 96-well microplate. Activity measured in the absence of NADH, as blank, was subtracted from the total activity. The PPDK activity was determined by coupling the formation of pyruvate from PEP to the formation of lactate from pyruvate via added lactic dehydrogenase, resulting in the oxidation of NADH (Olson et al., 2017). One unit (U) of PPDK enzyme activity was defined as the liberation of 1 μmol of NAD^+^. The protein concentration was determined using BioDrop (Indolab, Utama).

PPDK activity was calculated using following formula:

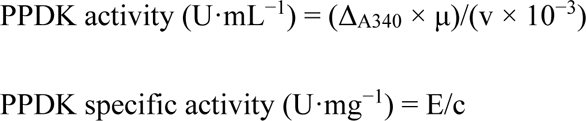

Here, μ is the dilution factor of the enzyme, v is the added amount of enzyme, E is the PPDK activity (U·mL^−1^), and c is the protein concentration (mg·mL^−1^).

### Western Blot Analysis

The crude extract samples were separated by SDS-PAGE on 10% gels as described previously (Mamedov et al., 2017), followed by transfer to Polyvinylidene fluoride (PVDF) membranes (Millipore, Billerica, MA) for western blot analysis. After transfer, the membranes were blocked using a 0.5% I-Block (Applied Biosystems, Carlsbad, CA). The membranes were first washed with a Phosphate-buffered saline, containing 0.1% Tween-20 (PBS-T). The His tagged proteins were probed with a purified mouse anti-His tag primary antibody (Cat. no. 652502, BioLegend) and anti-mouse horseradish peroxidase (HRP)-conjugated IgG secondary antibody (Cat. No. ab98790, Abcam). The protein bands in the blot wereusing a chemiluminescent substrate (SuperSignal West Pico, Thermo Fisher Scientific, Grand Island, NY).

### Protein sequence analyses and phylogenetic tree construction

Protein sequences of different PPDKs were obtained through “PPDK” or “pyruvate ortophosphate dikinase” keyword searches in Phytozome (Goodstein et al., 2012, https://phytozome-next.jgi.doe.gov/) and NCBI databases (https://www.ncbi.nlm.nih.gov/). Amino acid sequence alignments of *C. reinhardtii* PPDK1 and PPDK2 were performed using the MAFFT (Multiple Alignment Using Fast Fourier Transform, version 7, https://mafft.cbrc.jp/alignment/server/index.html) program with representative plant, algal, cyanobacterial PPDKs. MEGA–X (Molecular Evolutionary Genetics Analysis) software was used to align amino acid sequences from PPDK proteins. Multiple sequence alignments were constructed using ClustalW and MUSCLE algorithms implemented,using default settings. A phylogenetic tree was constructed using ETE3 (Huerta-Cepas et al., 2016). Domain predictions were made using Pfam (Mistry et al., 2021).

### Gene expression analysis by reverse transcription quantitative real-time PCR (RT-qPCR)

The samples prepared by first-strand cDNA synthesis were diluted 1:50. The experiment was carried out on a 96-well plate and each reaction contained 12.5 μL SYBR Green Master Mix (Thermo Scientific), 5 μL of cDNA, 0.3 μM each of the forward and reverse primers, and water to finally volume 25 μL. PCR was done in Roche Light Cycler 96. Reaction conditions were 10 min at 95°C, followed by 40 cycles of 95°C for 15 s, 60°C for 20 s, 72°C for 30 s, and a 1 s hold at 421 95°C to eliminate background fluorescence generated by the formation of primer dimers, followed by quantification of the fluorescence. The primer sequences are given in Table 5. Relative expression levels were normalized using the *C. reinhardtii* reference CBLP gene (Chang et al., 423 2005). 2 _-ΔCt_ statistical approach was performed for gene expression stability. Melt curve analyses were performed to determine whether the products represent a single specific species.

**Table 5.**
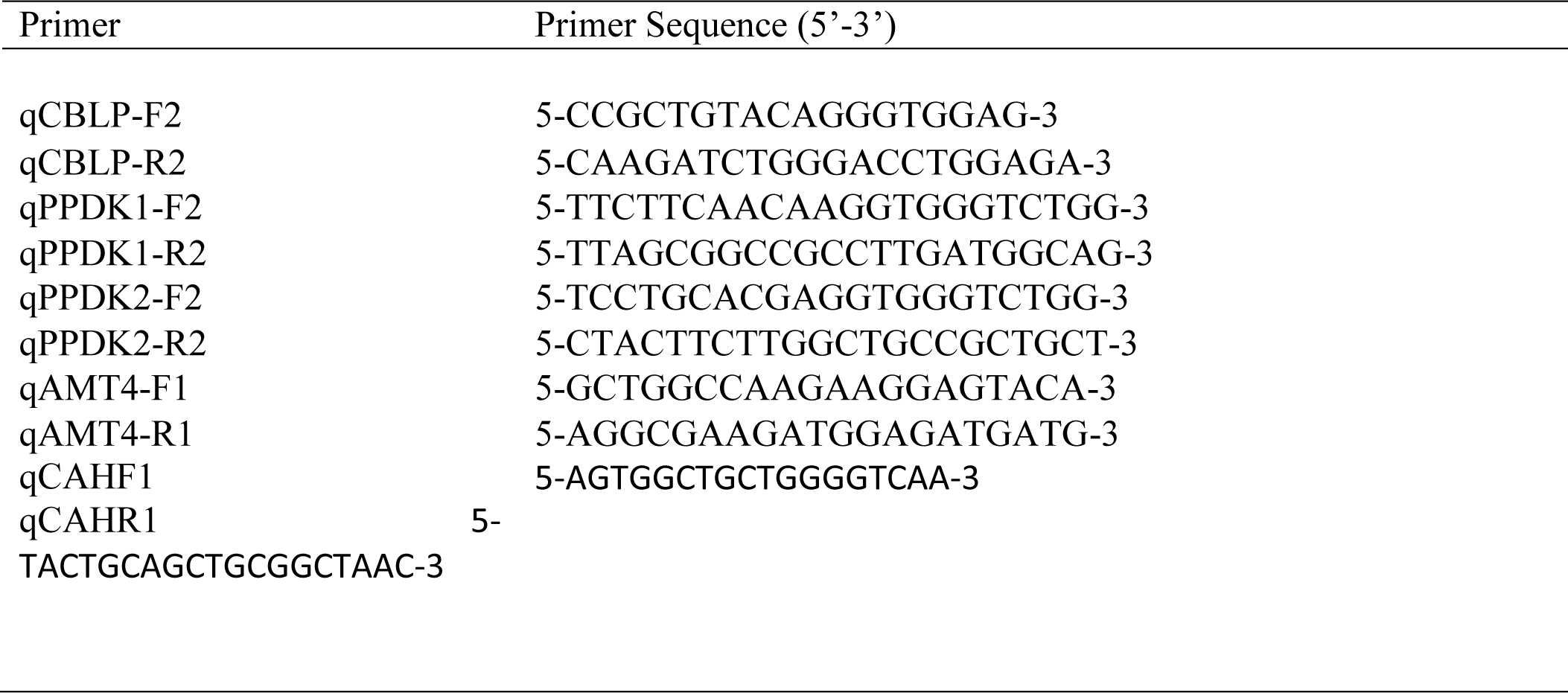
Primers used in qRT-PCR analysis.

The ammonium transporter (Amt) gene, which encodes a putative gas channel for the uncharged NH_3_ species (Soupene et al. 2004) was used as an N-sensitive reference gene (Mamedov et al. 2005; 2024). The carbonic anhydrase (CAH) gene was used as a CO_2_-sensitive reference gene (REFERENCE).

### Statistical analysis

For statistical analysis, the GraphPad Prism 9 program was used. *Student*’s t-*test* was used to evaluate the findings statistically. The results were evaluated within the 95% confidence interval and the significance level was p<0.05.

## CONCLUSIONS

Our long-term research goals are engineering *C. reinhardtii* metabolism to enhance the accumulation of TAGs while maintaining the efficiency of carbon dioxide conversion. The immediate goal of this study is a detailed molecular and biochemical characterization of PPDK genes for further knockout of these genes in *C. reinhardtii*. Our central hypothesis is that a detailed characterization of C4 metabolism in *C. reinhardtii* may lead to strategies for increasing the levels of TAGs, by modifying levels of storage oil/protein, keeping in mind that i) photosynthetic carbon flux tends to synthesize proteins or lipids depending on the relative activity and ii) lipids and proteins of cells compete for the same substrate, i.e., pyruvate, a product of glycolysis, and iii) the levels of pyruvate and TAGs can be significantly altered/regulated by C4-methabolism enzymes. Carbon skeletons generated through this pathway can contribute to the synthesis of acetyl-CoA, a central metabolite in lipid metabolism. Acetyl-CoA serves as a precursor for fatty acid synthesis, a process that ultimately leads to the production of TAGs. Therefore, regulating PPDK activity can indirectly influence the availability of acetyl-CoA for TAG synthesis. In addition, understanding the role of PPDKs and the molecular mechanism underlying the broader photosynthetic pathway in green algae including *C. reinhardtii* and other photosynthetic organisms would be useful for increasing the productivity of these organisms by optimizing photosynthetic efficiency that would be useful for applications such as biofuel production.

## Abbreviations

PPDK: pyruvate, ortophosphate dikinase
PEP: phosphoenolpyruvate
NADH /NAD^+^: Nicotinamide adenine dinucleotide

## Acknowledgments

Authors thank Dr. Gunay Mammadova for editorial assistance. All authors declare that they have no conflict of interest. This study was funded by the Scientific and Technological Research Council of Turkey (TUBITAK) (www.tubitak.gov.tr) through the project coded 214Z118 to Tarlan Mamedov.

## AUTHOR CONTRIBUTIONS

TM conceptualized this study, supervised the research, and wrote the manuscript. FD, FM, NC, TT, GM and MO performed experiments. FD and GH helped write the manuscript.

